# Potential for rapid genetic adaptation to warming in a Great Barrier Reef coral

**DOI:** 10.1101/114173

**Authors:** Mikhail V. Matz, Eric A. Treml, Galina V. Aglyamova, Madeleine J. H. van Oppen, Line K. Bay

**Affiliations:** Department of Integrative Biology, University of Texas at Austin, 205 W 24th St. C0990, Austin, Texas 78712, USA; School of BioSciences, University of Melbourne, Victoria 3010 Australia; Australian Institute of Marine Science, QLD, Australia

## Abstract

Can genetic adaptation in reef-building corals keep pace with the current rate of sea surface warming? Here we combine population genomic, biophysical modeling, and evolutionary simulations to predict future adaptation of the common coral *Acropora millepora* on the Great Barrier Reef. Loss of coral cover in recent decades did not yet have detectable effect on genetic diversity in our species. Genomic analysis of migration patterns closely matched the biophysical model of larval dispersal in favoring the spread of existing heat-tolerant alleles from lower to higher latitudes. Given these conditions we find that standing genetic variation could be sufficient to fuel rapid adaptation of *A. millepora* to warming for the next 100-200 years, although random thermal anomalies would drive increasingly severe mortality episodes. However, this adaptation will inevitably cease unless the warming is slowed down, since no realistic mutation rate could replenish adaptive genetic variation fast enough.

---

Hot water coral bleaching, caused by global warming, is devastating coral reefs around the world (*1*) but there is room for hope if corals can adapt to increasing temperatures (*2*). The fact that current coral generation is suffering high mortality does not necessarily imply that the next coral generation would not be better adapted. Many coral species have wide distributions that span environments that differ dramatically in their thermal regimes, demonstrating that efficient thermal adaptation has occurred in the past (*3*). But can coral adaptation keep up with the unprecedentedly rapid current rate of global warming (*4*)? One way for corals to achieve rapid thermal adaptation is through genetic rescue, involving the spread of existing heat tolerance alleles from warm-adapted populations to now-warming regions via larval migration (*5*, *6*). We have previously demonstrated the presence of genetic variants conferring high thermal tolerance in a low-latitude *A. millepora* population (*5*). It can be hypothesized that global warming will cause preferential survival of migrants from warmer to cooler locations because they will be following their thermal optimum, whereas individuals migrating in the opposite direction would find themselves in increasingly mismatched environments (Fig. 1 A, B). Another likely population-level effect of recent declines in coral cover (*7*) is a reduction in overall genetic diversity, potentially limiting both the scope and the rate of adaptation. Here, we tested these predictions in *Acropora millepora*, a common reef-building coral from the most ecologically prominent and diverse coral genus in the Indo-Pacific (staghorn corals, *Acropora*), and used the obtained demographic estimates to model the future adaptive potential of *A. millepora* on the Great Barrier Reef (GBR).

**Figure 1.**
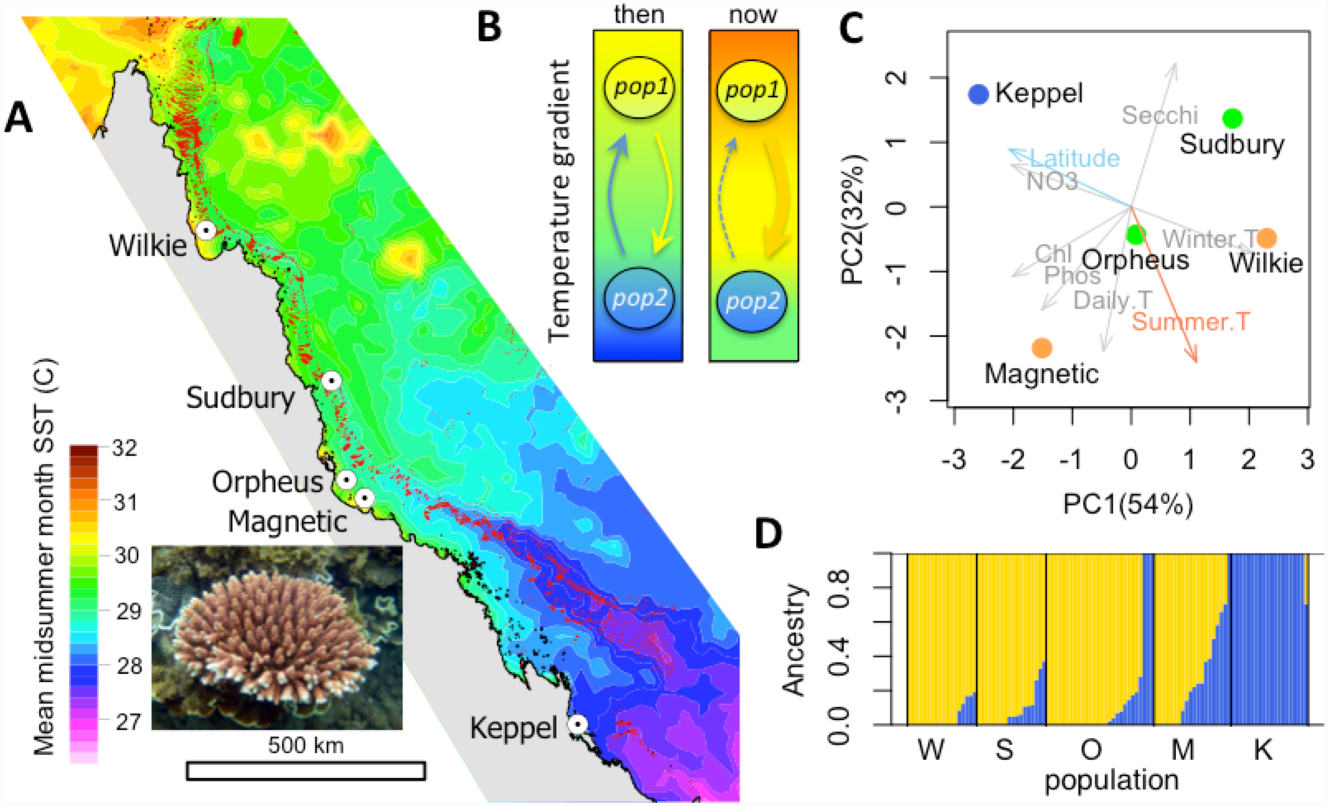
The population setting and background for our study. (A) Locations of sampled populations where mean midsummer month sea surface temperature differed by up to ~3°C. Inset: *Acropora millepora*. (B) Hypothesized migration change under global warming: Warm-adapted genotypes that migrate to locations that used to be cooler would be following their physiological optimum and hence expected to survive better than migrants in the opposite direction. (C) Principal component analysis of water quality and temperature parameters at the sampled locations. Winter.T - 10% quantile of winter temperature, Summer.T – 90% quantile of summer temperature, Daily.T – 90% quantile of daily temperature range, Phos – total dissolved phosphorus, Chl – chlorophyll, NO3 – nitrate, Secchi – Secchi depth (water clarity). Locations are colored according to summer temperature. (D) ADMIXTURE plot of ancestry proportions with *K* = 2.

## Locations and genotyping

We used samples collected in 2002-2009 from five populations of *A. millepora* along the latitudinal range of the GBR (Fig. 1 A). Environmental parameters (obtained from <u>http://eatlas.org.au/</u>) varied widely among these locations (Fig. 1 C). Importantly, maximum summer temperature (the major cause of bleaching-related mortality) followed the latitudinal gradient with one notable exception: one of the near-shore populations from the central GBR (Magnetic Island) experienced summers as hot as the lowest-latitude population (Wilkie Island (Fig. 1 C).

We genotyped 18-28 individuals per population using 2bRAD (*8*) at >98% accuracy and with a >95% genotyping rate. Analysis of population structure based on ~11,500 biallelic SNPs agreed with previous results (*9*, *10*) and revealed very low levels of genetic divergence, with only the Keppel Islands population being potentially different from the others (Fig. 1 D and Fig. S1). Pairwise *F*_ST_ were small and did not exceed 0.014 even between the southernmost and northernmost populations (Keppel and Wilkie).

## Demographic subdivision and migration patterns

To more rigorously test for population subdivision and infer unidirectional migration rates among populations and population sizes, we used Diffusion Approximation for Demographic Inference (*dadi*, (*11*)). *dadi* is a method that optimizes parameters of a pre-specified demographic model to maximize the likelihood of generating the observed allele frequency spectrum (AFS). For two populations AFS is essentially a two-dimensional histogram of allele frequencies (Fig. S2). Being a likelihood-based method, *dadi* can be used to compare alternative models using likelihood ratio tests and Akaike Information Criterion (AIC). Most importantly, unlike previously used approaches (*10*, *12*, *13*) *dadi* does not rely on assumptions of genetic equilibrium (stability of population sizes and migration rates for thousands of generations) or equality of population sizes and therefore is potentially more realistic and sensitive for natural populations.

We used bootstrap-AIC approach to confirm that our populations are separate demographic units. For each pair of populations we generated 120 bootstrapped datasets by resampling genomic contigs and performed delta-AIC comparison of two demographic models, a split-with-migration model and a no-split model (Fig. S3 C). The split-with-migration model assumed two populations that split some time *T* in the past with potentially different sizes *N1* and *N2*, and exchange migrants at different rates (*m12* and *m21*) depending on direction. The no-split model allowed for ancestral population size to change but not for a population split, so the experimental data were modeled as two random samples from the same population of size *N*. The majority of bootstrap replicates (64-100%) showed AIC advantage of the split-with-migration model for all pairs of populations except Sudbury-Magnetic (41% support, Fig. S3). This indicates that most *A. millepora* populations on the GBR are in fact demographically distinct, despite typically non-significant *F*_ST_ reported by previous studies based on allozymes (*12*, *13*) and microsatellite markers (*10*).

AFS-based analysis allows rigorous estimation of unidirectional migration rates between populations. The classical *F*_ST_ – based approach only allows estimating bi-directional migration rate (*13*) and even that calculation has been criticized because its underlying assumptions are rarely realistic (*14*). We determined unidirectional migration rates from the split-with-migration model and estimated their confidence limits from bootstrap replicates. In theory, migration rate can be confounded with population divergence time, since in the AFS higher migration often looks similar to more recent divergence (*15*). To confirm that the model with ancient population divergence and migration is preferable to the model with very recent divergence and no migration, we performed the delta-AIC bootstrap comparison between these models and obtained overwhelming support for the model with ancient divergence and migration (Fig. S4). Notably, for all pairwise analyses migration in southward direction exceeded northward migration, and this difference was significant in seven out of nine cases (Fig. 2 A and Fig. S3A, C). Linear mixed model analysis of direction dependent median migration rates with a random effect of destination (to account for variation in total immigration rate) confirmed the overall significance of this southward trend (*P*_MCMC_ <1e-4). Full listing of parameter estimates and their bootstrap-derived 95% confidence limits is given in Table S1.

**Figure 2.**
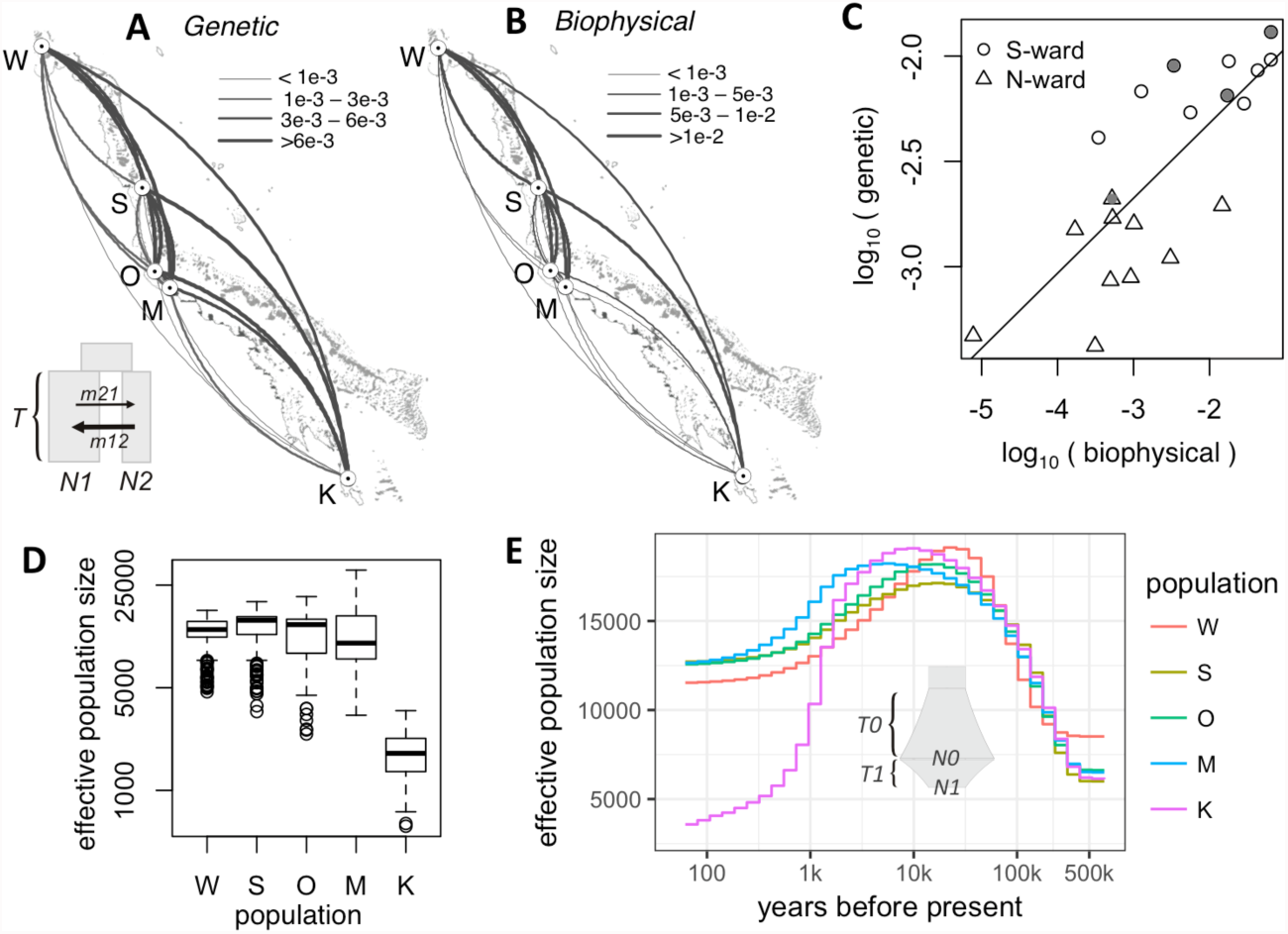
Demography of *A. millepora* populations on the GBR. (A) Arc-plot of migration rates among populations reconstructed from population genetic data. Inset: *dadi* model used: ancestral population splits into two populations of unequal sizes (N1 and N2) some time T in the past, these populations exchange migrants at different rates depending on direction. (B) Migration rates according to the biophysical model. On panels A and B, the arcs should be read clockwise to tell the direction of migration; line thickness is proportional to the migration rate. (C) Correlation between log-transformed biophysical and genetic migration rates (Mantel r = 0.58, *P* = 0.05). Grey symbols are migration rates to Magnetic Island, the high-temperature location in the central GBR. (D) Box plot of effective population sizes inferred by the split-with-migration model (panel A) across all population pairs and bootstrap replicates. (E) Historical changes in effective population sizes inferred using a single-population model with two periods of exponential growth (*T0* and *T1*, reaching sizes *N0* and *N1*, inset), averaged across bootstrap replicates.

To investigate whether the southward migration bias was due to higher survival of warm-adapted migrants, as predicted under global warming (Fig. 1 B), we developed a biophysical model of coral larval dispersal on the Great Barrier Reef. This model quantified the per-generation migration potential among coral reef habitat patches in the GBR based on ocean currents and parameters of larval biology (*16*, *17*). The biophysical model predicted very similar migration rates as our genetic model (Mantel *r* =0.58, *p* = 0.05), recapitulating the southward bias (Fig. 2 A-C). Importantly, the same southward bias was predicted for population pairs in which southward migration corresponded to movement to the same-temperature or even to warmer location, such as migrations to the Magnetic Island (Fig. 2 C, grey points). This indicates that southward migration bias is predominantly driven by ocean currents and not by preferential survival of warm-adapted coral genotypes migrating to cooler locations.

Migration estimates from our main genetic model (Fig. 2 A, inset) represented historical averages since the populations split and did not resolve any potential recent migration changes. To determine if there were any recent changes in southward migration, we evaluated an extended split-with-migration model that allowed for a change in migration over the past 75-100 years. The extended model suggested some recent migration changes, including southward migration increases (Fig. S5) but once again, these changes did not correspond with an increase in migration from warmer to cooler locations. We conclude that with the current data and analysis techniques we cannot (yet) detect an effect of recent warming on preferential direction of coral migration on the GBR.

## Genetic diversity trends

The GBR has warmed considerably since the end of last century (*18*), which may have already reduced genetic diversity in *A. millepora* populations. We used *dadi* to infer effective population sizes, which is a measure of genetic diversity and one of the key parameters determining the population’s adaptive potential (*19*). The results of the split-with-migration model (Fig. 2 A) were consistent for all population pairs and indicated that Keppel population was about one-fifth the size of others (Fig. 2 D, E). This result was not surprising since the Keppel population frequently suffers high mortality due to environmental disturbances (*9*). We also used a single-population *dadi* model that allowed for two consecutive growth/decline periods (Fig. 2 E, inset) to reconstruct effective sizes of individual populations through time (Fig. 2 E and Fig S6). All populations showed evidence of growth prior to the last glaciation, between 500 and 20 thousand years ago (Fig 2 E), which aligned well with the fossil record of rising dominance of *Acropora* corals on Indo-Pacific reefs during this period (*20*). It was hypothesized that the fast growth and early sexual maturation of *Acropora* corals gave them an advantage relative to most other reef-building corals during dynamic changes in the reef-forming zone due to the sea level changes accompanying glacial cycles (*20*). Our results suggest that *A. millepora* populations have been in decline since sea level stabilized after the last deglaciation, roughly 10 thousand years ago (Fig. S6). This decline was the most pronounced for the Keppel population (Fig. 2 E), but delta-AIC bootstrap supported inclusion of additional growth/decline period for all populations except Magnetic Island (Fig. S7). None of the populations showed evidence of accelerated decline in effective population size over the past hundred years, despite recent GBR-wide decline in overall coral cover (*7*). It must be noted, however, that recent decline in population size is hard to detect using the AFS method unless the sample size is very large, since it would predominantly affect the frequencies of rare alleles (*15*, *21*). Despite this shortcoming, we can conclude that genetic diversity in *A. millepora* has not yet been strongly affected by warming over the past century, although the populations appear to have been in long-term decline GBR-wide for the past several thousand years.

## Metapopulation adaptation model

To evaluate whether standing genetic variation contributed by local thermal adaptation could facilitate rapid adaptation of the *A. millepora* metapopulation in response to warming, we developed an individual-based multigene model of metapopulation adaptation in the SLiM software environment (*22*). The model’s code is highly flexible and can simulate any number of populations with any configuration of population sizes, migration rates, and environmental trends. The model also allows varying the number and effect sizes of QTLs, heritability, and phenotypic plasticity. Here, we used population sizes and migration rates inferred from the genetic analysis (Fig. 2 A, D) and incorporated differences in midsummer monthly mean temperature among populations (Fig. 1 A). The populations were allowed to adapt to local thermal conditions for 2,000 generations. Assuming a generation time of 5 years in *A. millepora* (*23*) this corresponded to the period of stable temperature since the last deglaciation. After this pre-adaptation, the temperature began to increase at a rate of 0.05°C per generation in all populations, corresponding to the projected 0.1°C warming per decade (*24*). During warming period, a population declining in fitness would shrink in size and stop contributing migrants to other populations. Throughout the simulation the temperature was allowed to fluctuate randomly between generations to approximate El Nino Southern Oscillation (ENSO).

We found that, with only ten thermal QTLs, under a broad range of settings for heritability and plasticity the pre-adapted metapopulation was able to persist through the warming for at least 20 - 50 generations (100 - 250 years) and, under some parameter combinations, much longer (Fig. 3 and S8). Migration substantially contributed to this persistence (Fig. 3 E, F), underscoring the importance of divergent local adaptation and genetic rescue (*5*, *6*). However, when existing standing genetic variation was finally depleted, all populations inevitably went extinct beginning with warmer locations. Allowing for new mutations could not avert this extinction even when the mutation rate was extremely high - 1e-5 per QTL per gamete or higher while assuming that half of all mutations were beneficial (Fig. 4).

**Figure 3.**
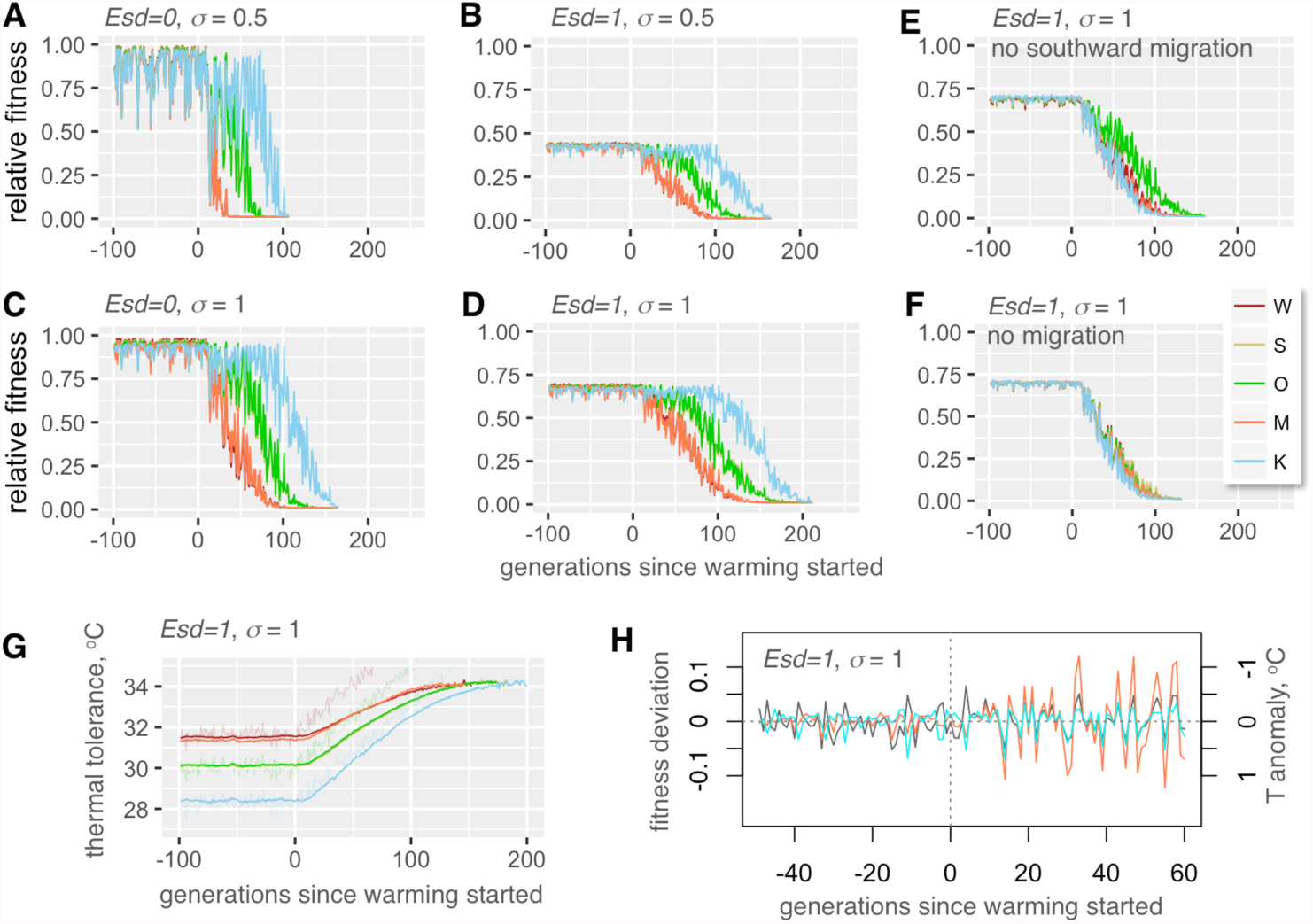
Modeling coral metapopulation persistence under warming based on standing genetic variation (no new mutations). (A-D) Fitness of modeled populations depending on the magnitude of non-heritable phenotypic component (*Esd*, standard deviation of normally distributed random value added to the sum of QTL effects, in degrees C), phenotypic plasticity (σ, standard deviation of the Gaussian slope of fitness decline away from the phenotypic optimum, in degrees C), and presence-absence of migration (E, F). On panels A-F, y-axis is observed fitness relative to maximum attainable with perfect heritability at the genetically determined optimum, averaged over all individuals in a population. Warm-adapted populations (W and M) are shown as red-tint traces, populations from mild thermal regime (S and O) are green-tint traces, and the cool-adapted population (K) is the blue trace. Note nearly complete overlap between traces for pairs of populations pre-adapted to the same temperature (W, M and S, O). (G) Thermal tolerances of evolving populations. Thin noisy lines are modeled temperatures at different locations. (H) Modeled random temperature anomalies (grey line) and fluctuations in populations’ fitness (colored lines: residuals from loess regression over fitness traces on panel D; Wilkie: orange line, Keppel: blue line). Note the inverse sign of temperature anomalies: this more clearly shows the correspondence between rise in temperature and drop in fitness. As warming progresses, populations (especially originally warm-adapted ones) become increasingly sensitive to random temperature fluctuations.

**Figure 4.**
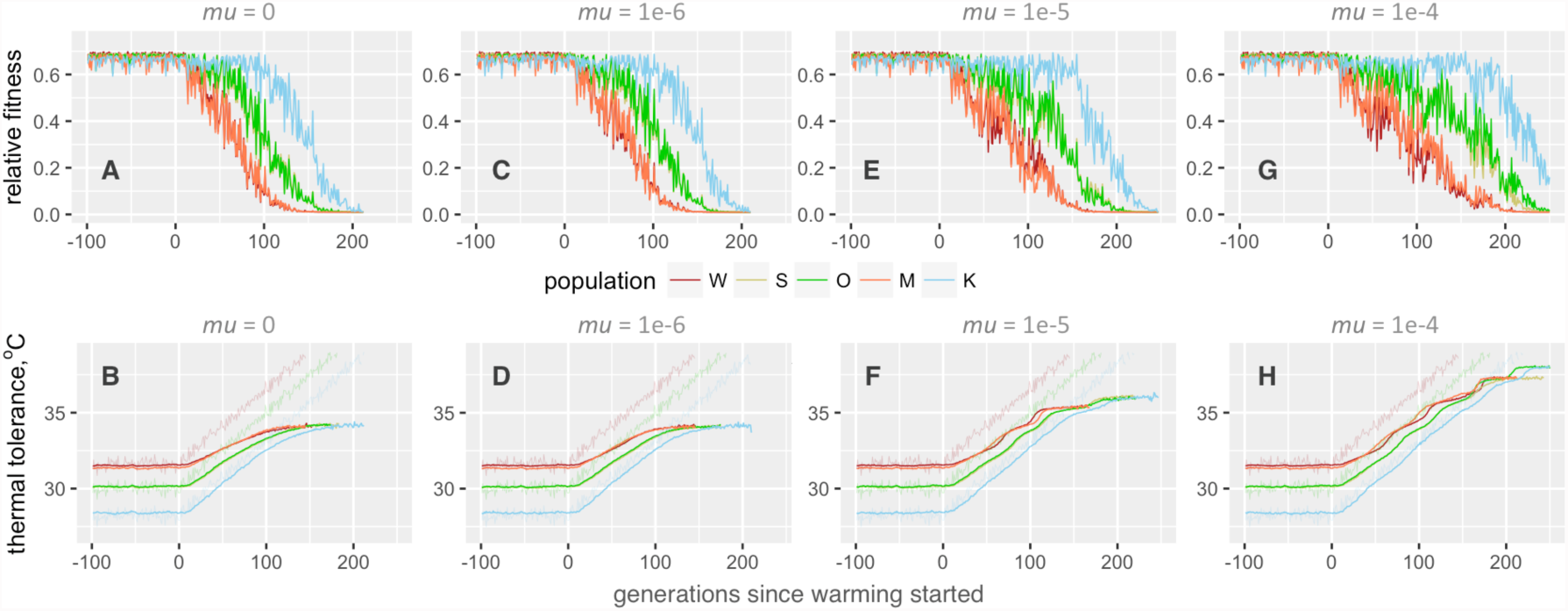
New mutations cannot prevent extinction. Top row of graphs (A, C, E and G) show population fitness; bottom row (B, D, F, H) – mean thermal tolerance. All cases share the settings of *Esd* = 1 and σ =1. Mutation rate (*mu*) per QTL per gamete is listed above the graphs; effect sizes of new mutations were drawn from a normal distribution with mean 0 and standard deviation 0.2. Note a few “evolutionary rescue” events at high mutation rates, when a new adaptive mutation spreads through metapopulation leading to temporary acceleration of phenotypic evolution (F, H).

A notable tendency observed with all parameter settings was that during warming the fitness (and hence the size) of adapting populations began to fluctuate following random thermal anomalies, and the amplitude of these fitness fluctuations increased as the warming progressed even though the amplitude of thermal anomalies did not change (Fig. 3 H). These fluctuations correspond to severe mortality events induced by thermal extremes that can occur as a result of ENSO and affected warm-adapted populations most, which very much resembles the situation currently observed throughout the world (*1*).

## What helps corals adapt?

Predictably, higher phenotypic plasticity promoted metapopulation persistence and stability against random thermal anomalies, but we were rather surprised to observe a similar positive effect of lower heritability (i.e., higher non-heritable component, Fig 3, S8 and S9). Low heritability of thermal tolerance is expected for reef-building corals: much of natural variation in this trait in corals is due to the type of algal symbionts (*Symbiodinium* spp. (*25*)). Photo-symbionts are not transmitted from parent to offspring in the majority of coral species (*26*), and although host genetics can have some effect on the choice of *Symbiodinium* in the next generation(*27*) environment has stronger effect on this association (*25*, *28*).

Longer metapopulation persistence under low heritability and high plasticity was most likely due to their enhancing effect on standing genetic variation (Fig. 5). Higher plasticity promoted both higher number of variants retained in populations and larger effect sizes of these variants, whereas higher heritability also led to higher number of retained variants but notably smaller effect sizes (Fig. 5 A-K). It can be said that under high heritability local adaptation was based on many mutations of small effect, whereas low heritability promoted adaptation involving fewer mutations of larger effect. Importantly, the cumulative absolute effect of QTL variants in a population was consistently higher under the setting of low heritability, despite lower number of variants (Fig. 5 K, L). During warming, this variation lasted longer as the source of adaptive genetic variants, enabling up to 4°C increase in mean thermal tolerance (Fig. 3 G and S9). Another important effect of higher plasticity was that it partially rescued the drop in average population fitness due to low heritability (Fig. 3 B, D and S9). This drop happened because low heritability prevented individuals from attaining maximum fitness even if their genetics was perfectly matched to the environment.

**Figure 5.**
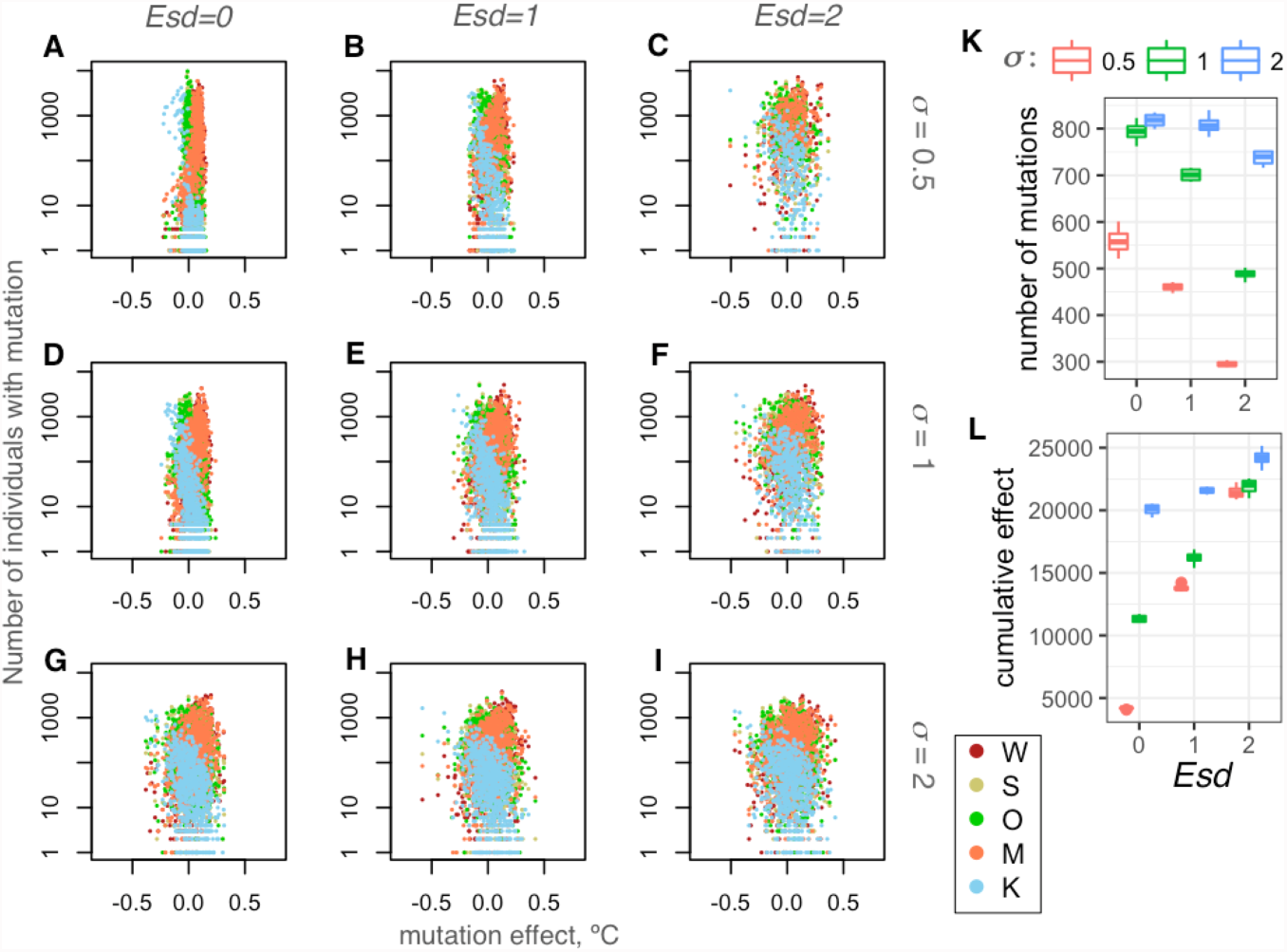
Effects of non-heritable phenotypic component (*Esd*) and phenotypic plasticity (σ) on standing genetic variation at thermal QTL loci. (A - I) Dependence of the number of individuals in each population bearing a mutation on the mutation’s effect size (change in thermal tolerance, in °C). Each mutation is represented by up to five points colored according to the population in which it is found (see legend). Lower heritability (higher *Esd*) and higher plasticity promote retention of mutations with larger effect sizes. (A, D, G): *Esd*=0. (B, E, H): *Esd*=1. (C, F, I): *Esd*=2. (A-C): σ = 0.5. (D-F): σ = 1. (G-I): σ = 2. (K, L) Genetic variation retained after 2000 generations of adaptation to benign local thermal conditions (aggregating 12 simulation replicates for two populations, S and O). (K) Number of mutations at QTL loci.(L) Cumulative effect size (sum of products of mutation’s absolute effect size and the number of individuals bearing the mutation). Lower heritability (higher *Esd*) results in retaining fewer mutations, but these mutations amount to up to four-fold larger cumulative effect size, while higher plasticity promotes both higher number of mutations and larger effect sizes.

## Potential pitfalls

There are several uncertainties in our model associated with coral biology. Below we argue that, while more research is certainly needed to resolve these uncertainties, our modeling was conservative overall.

We assumed only ten QTLs, which is likely much fewer that the actual number of thermal QTLs in acroporid corals (*20*). Higher number of QTLs and/or their larger effect sizes would promote higher genetic variation and lead to longer population persistence. We also kept the distribution of QTL effect sizes narrow: with the current settings and ten QTLs, at the start of simulation only about 2% of corals deviated from the mean thermal tolerance by more than 1.5°C in either direction. Such narrow variation makes adaptation to the thermal gradient of ~3°C along the GBR non-trivial, but still, at present there is no experimental data to evaluate whether even such narrow variation is realistic.

We used effective population sizes suggested by genetic analysis as census sizes. In highly fecund marine organisms census sizes tend to substantially exceed effective population sizes, sometimes by orders of magnitude (*29*), which would strongly promote higher genetic variation and population persistence. Moreover, we modeled only our five populations rather than the whole GBR, which would have resulted in much higher standing genetic variation in the metapopulation, promoting longer persistence.

As for phenotypic plasticity, in simulations shown on Figs. 3 σ = 0.5 and σ = 1 corresponded to 86% and 40% decline in fitness if the individual’s phenotype mismatched the environment by 1°C. The existing data on coral thermal plasticity are somewhat conflicting. One study shows that acroporid corals can successfully acclimatize to environments differing in maximum temperatures by as much as 2°C (*30*); however, another study found that coral grew 52-80% more slowly when transplanted among locations differing by 1.5°C average temperature, (*31*). Although it is not possible to directly interpret these results in terms of width of the fitness function (as plasticity is encoded in our model) the former study likely supports the higher plasticity setting (σ = 1) while the latter study supports σ = 0.5. It must also be noted that both these studies involved *in situ* transplantations and hence the effect of temperature remains confounded with other local fitness-affecting environmental parameters. Also, in adult corals plasticity is likely lower that in larvae and recruits, which are expected to exhibit non-reversible developmental plasticity associated with metamorphosis and establishment within a novel environment (*32*). One particularly important event during this developmental transition is establishment of association with local algal symbionts. Since symbionts also adapt to local thermal conditions (*28*) this would elevate the fitness of the coral host despite possible mismatch between its own genetically determined thermal optimum and local temperature, which in our model implies broadening of the fitness function (i.e., higher plasticity). Future experiments that expose multiple genetically distinct coral individuals to a range of temperatures under controlled laboratory settings are required to rigorously quantify variation in thermal optima and plasticity in natural populations.

Our demographic analysis and, by implication, our modeling results are contingent on the accuracy of the mutation rate estimate. The mutation rate used here, 4e-9 per base per year, or 10 mutations per genome per generation (see Methods), is but a rough estimate based on phylogenetic distances (*33*). Higher mutation rate would lead to smaller population sizes, higher migration rates, and more recent time stamps for population splits or population size changes (see Methods, Unit Conversion section). As a consequence, it would lead to lower standing genetic variation in the metapopulation and therefore to diminished potential to persist under global warming. Although the rate used here is within the ballpark of mutation rates measured in multicellular animals (*34*) in the future a more precise estimate of mutation rate should be obtained to sharpen the demographic estimates and parameterize the adaptation model more realistically.

It may be argued that our samples are genetically out of date, not capturing the effects of disturbances that happened on the GBR since the time of their collection (2002-2009). However, as we mentioned above, very recent demographic events (in our case, 2-3 generations ago) are undetectable at the level of neutral genetic variation unless the study’s sample size is comparable to the number of disturbance-surviving individuals (i.e., either when the disturbance was truly catastrophic or the sample size is very large). Thus, our samples can still be considered well representative of major patterns of genetic diversity of our study species.

Finally, our model assumed that recovery from high mortality events would happen without impediment, through reseeding by survivors and migrant influx from other coral populations. However, ecological feedbacks such as shifts to an alternative ecological stable state (*35*) might substantially decrease the rate of reseeding and recovery of affected reefs. In that case, the increase in severity of bleaching-related mortality might lead to much faster coral extinction than predicted by our model.

## Conclusions

Our study provides a novel integrated empirical and modeling framework to evaluate the risk of extinction in natural populations. We found that genetic diversity and migration patterns of *Acropora millepora* were not yet strongly affected by climate change and were well positioned to facilitate persistence of the GBR metapopulation for a century or more. Our results underscore the pivotal role of standing genetic variation and genetic exchange in the future metapopulation persistence. This implies that any intervention that would reduce this variation (for example, captive breeding based on selection of only a few “winner” genotypes or clonally propagating a small number of genotypes for reef restoration) is likely to have negative impact on corals’ adaptive potential. In contrast, efforts facilitating the spread of genetic variation, such as assisted gene flow (*36*), could be much more helpful in the long term. With the estimated natural migration rates on the order of 5-100 migrants per generation, human-assisted genotype exchange could appreciably contribute to the genetic rescue without risking disruption of the natural local adaptation patterns (*37*). However, despite good prospects for short-term adaptation, corals are predicted to become increasingly more sensitive to random thermal anomalies, especially in the originally warm-adapted populations. The 10-85% mortality in the Northern GBR as a result of 2016 bleaching event (*38*) could be a particularly sobering recent manifestation of this trend. Finally, it is important to point out that adaptation based on genetic rescue will not save corals from eventual extinction: it will only buy us some time to take action against global warming, which hopefully can be stopped before corals run out of genetic variation.

## Methods

### Genotyping

This study relied predominantly on samples described by van Oppen et al (*10*), with addition of several samples from Orpheus and Keppel islands that were used in the reciprocal transplantation experiment described by Dixon et al (*39*). The samples were genotyped using 2bRAD (*8*) modified for Illumina sequencing platform; the latest laboratory and bioinformatics protocols are available at <u>https://github.com/z0on/2bRAD_GATK</u>. BcgI restriction enzyme was used and the samples retained for this analysis had 2.3-20.2 (median: 7.45) million reads after trimming and quality filtering (no duplicate removal was yet implemented in this 2bRAD version). The reads were mapped to the genome of the outgroup species, *Acropora digitifera* (*40*, *41*), to polarize the allelic states into ancestral (as in *A. digitifera*) and derived, e.g., (*42*, *43*). Genotypes were called using GATK pipeline (*44*).

Preliminary analysis of sample relatedness using vcftools (*45*) revealed that our samples included several clones: four repeats of the same genotype from the Keppel Island (van Oppen et al (*10*) samples K210, K212, K213 and K216), another duplicated genotype from Keppel (samples K211 and K219), and one duplicated genotype from Magnetic Island (samples M16 and M17). All other samples were unrelated. We took advantage of these clonal replicates to extract SNPs that were genotyped with 100% reproducibility across replicates and, in addition, appeared as heterozygotes in at least two replicate pairs (script replicatesMatch.pl with hetPairs=2 option). These 7,904 SNPs were used as “true” SNP dataset to train the error model to recalibrate variant quality scores at the last stage of the GATK pipeline. During recalibration, we used the transition-transversion (Ts/Tv) ratio of 1.438 determined from the “true” SNPs to assess the number of false positives at each filtering threshold (as it is expected that an increase of false positive calls would decrease the Ts/Tv ratio towards unity). We chose the 95% tranche, with novel Ts/Tv = 1.451. After quality filtering that restricted the calls to only bi-allelic polymorphic sites, retained only loci genotyped in 95% or more of all individuals, and removed loci with the fraction of heterozygotes exceeding 0.6 (possible lumped paralogs), we ended up with 25,090 SNPs. In total, 2bRAD tags interrogated 0.18% of the genome. The genotyping accuracy was assessed based on the match between genotyped replicates using script repMatchStats.pl. Overall agreement between replicates was 98.7% or better with the heterozygote discovery rate (fraction of matching heterozygote calls among replicates) exceeding 96%.

### Genome-wide genetic divergence

To begin to characterize genome-wide divergence between populations we used pairwise genome-wide Weir and Cockerham’s *F*_ST_ calculated by vcftools (*45*), principal component analysis (PCA) using R package adegenet (*46*), and ADMIXTURE (*47*). For PCA and ADMIXTURE, the data were thinned to keep SNPs separated by 5kb on average and by at least 2.5 kb, choosing SNPs with highest minor allele frequency (script thinner.pl with options ‘interval=5000 criterion=maxAF’). The optimal K in ADMIXTURE analysis was determined based on the cross-validation procedure incorporated within ADMIXTURE software; the lowest standard error in cross-validation was observed at K=1.

### Demographic analysis and bootstrapping

Prior to demographic analysis, Bayescan (*48*) was used to identify sites potentially under selection among populations, and 73 sites with q-value <0.5 were removed. This aggressive removal of potential non-neutral sites resulted in better agreement between bootstrap replicates compared to an earlier of analysis where only 13 sites with q-value < 0.05 were removed. Demographic models were fitted to 120 bootstrapped datasets, which were generated in two stages. First, three alternatively thinned datasets were generated for which SNPs were randomly drawn to be on average 5 kb apart and not closer than 2.5 kb. This time the SNPs were drawn at random to avoid distorting the allele frequency spectrum (unlike thinning for PCA and ADMIXTURE where the highest minor allele frequency SNPs were selected). Then, 40 bootstrapped replicates were generated for each thinned dataset by resampling contigs of the reference genome with replacement (script dadiBoot.pl). The fitted model parameters were summarized after excluding bootstrap replicates that fell into the lowest 15% likelihood quantile and the ones where model fitting failed to converge, leading to some parameters being undetermined or at infinity (less than 10% of total number of runs). Delta-AIC values were calculated for each bootstrap replicate that passed these criteria for both compared models, and summarized to obtain bootstrap support value, the percentage of replicates favoring the alternative model. While fitting *dadi* models, the data for each population were projected to sample sizes maximizing the number of segregating sites in the analysis, resulting in 7000-8172 segregating sites per population. Initially, our models included a parameter designed to account for ancestral state misidentification rate when constructing the polarized AFS (e.g., (*49*)), but since this parameter was consistently estimated to be on the order of 0.001 and had negligible effect on the models’ likelihood, we removed it from the final set of models.

### Unit conversion

To convert *dadi*-reported coalescent parameter values (*θ*, *T* and *M*) into time in years (*t*), effective population sizes in number of individuals (*Ne*) and migration rates as fraction of new immigrants per generation (*m*), we estimated the mutation rate (*μ*) from the time-resolved phylogeny of *Acorpora* genus based on *paxC* intron (*33*), at 4e-9 per base per year. Although *A. millepora* can reproduce after 3 years (*23*) we assumed a generation time of 5 years reasoning that it would better reflect the attainment of full reproductive potential as the colony grows. Assuming a genome size of 5e+8 bases (*40*) the number of new mutations per genome per generation is 10. Since the fraction 2bRAD-sequenced genome in our experiment was 1.8e-3, the mutation rate per 2bRAD-sequenced genome fraction per generation is *μ* = 0.018. This value was used to obtain:

- Ancestral effective population size: *Ne* = *θ* / 4*μ*
- Migration rate: *m* = *M* / 2*Ne*
- Time in years: *t* = 2*TNe* • 5

### Biophysical model

A spatially-explicit biophysical modeling framework (*16*, *50*) was used to quantify migration between coral reef habitats of the broader region surrounding the Great Barrier Reef, thereby revealing the location, strength, and structure of a species’ potential population connectivity. The model’s spatial resolution of ca. 8 km coincides with hydrodynamic data for the broader region (1/12.5 degree; HYCOM+NCODA Reanalysis and Analysis product; hycom.org). Our biophysical dispersal model relies on geographic data describing the seascape environment and biological parameters capturing coral-specific life-histories. Coral reef habitat data are available from the UNEP World Conservation Monitoring Centre (UNEP-WCMC; http://data.unep-wcmc.org/datasets/1) representing a globally-consistent and up-to-date representation of coral reef habitat. To capture specific inter-annual variability, two decades of hydrodynamic data were used from 1992 to 2013 (*51*).

Coral-specific biological parameters for *A. millipora* included relative adult density (dependent on the habitat), reproductive output, larval spawning time and periodicity (e.g., Magnetic Island populations spawn a month earlier than the other GBR sites (*52*)), maximum dispersal duration, pre-competency and competency periods, and larval mortality (*53*, *54*). The spatially explicit dispersal simulations model the dispersal kernel (2-D surface) as a ‘cloud’ of larvae, allowing it to be concentrated and/or dispersed as defined by the biophysical parameters. An advection transport algorithm is used for moving larvae within the flow fields (*55*).

Simulations were carried out by releasing a cloud of larvae into the model seascape at all individual coral reef habitat patches and allowing the larvae to be transported by the currents. Ocean current velocities, turbulent diffusion, and larval behavior move the larvae through the seascape at each time-step. Larval competency, behavior, density, and mortality determine when and what proportion of larvae settle in habitat cells at each time step. When larvae encounter habitat, the concentration of larvae settling with the habitat is recorded at that time-step. From the dispersal data, we derived the coral migration matrix representing the proportion of settlers to each destination patch that came from a source patch, which is analogous to the source distribution matrix (*56*) and is equivalent to migration matrices derived from population genetic analysis. It is important to note that migration matrices extracted for the field sites represent the potential migration through all possible stepping-stones.

### Metapopulation adaptation model

The model was implemented in SLiM (*22*), the forward evolutionary simulator, by modifying the provided recipe “Quantitative genetics and phenotypically-based fitness”. The model simulates Fisher-Wright populations with discreet generations. At the start of the simulation, populations are established at specified population sizes and pairwise migration rates (genetic replacement rates), and all QTLs in all individuals are given a mutation with the effect size drawn from a normal distribution with mean zero and specified standard deviation, to create standing genetic variation. The phenotype of each individual is calculated as the sum of QTL effects plus random noise of specified magnitude to simulate non-heritable phenotypic component. Then, fitness of each individual is calculated based on the difference between the individual’s phenotype (thermal optimum), temperature of the environment, and the setting for phenotypic plasticity, modeled as the width of the fitness curve: the standard deviation of the Gaussian slope of fitness decline away from the phenotypic optimum. Then, parents are chosen to produce the next generation according to their fitness; parents for immigrant individuals are chosen from among individuals in the source population. New mutations at QTLs happen at the specified rate when transitioning to the next generation and the effect of a new mutation adds to the previous QTL effect. To better model population dynamics, we implemented linear scaling of the population size and immigration rates with the population’s mean fitness. In the model described here this scaling was applied during warming period, so that a population declining in fitness relative to its state at the end of pre-adaptation period shrinks in size and stops contributing migrants to other populations.

Adjustable model parameters and their settings in this study:

- Number of QTLs and the distribution of their effect sizes. To keep the model conservative, we modeled only ten QTLs with normal distribution of effect sizes with a standard deviation of 0.2°C. With ten QTLs, this setting implied that at the start of simulation only about 2% of corals deviated from mean thermal tolerance by more than 1.5°C in either direction. Since thermal differences between our populations exceeded 3°C, this narrow variation made local adaptation rather non-trivial.
- Dominance of QTLs (set to 0.5 in our simulation).
- Phenotypic plasticity: standard deviation of the Gaussian curve describing decline in fitness away from phenotypic optimum. We modeled three plasticity settings, 0.5, 1 and 2, which corresponded to 86%, 40% and 13% fitness drop when the environment mismatched phenotypic optimum by 1°C.
- Non-heritable phenotypic component: standard deviation of a normal distribution with mean zero from which a random value is drawn to be added to the sum of QTL effects when computing phenotype. Setting this parameter to zero corresponds to trait heritability of one. Higher values of this parameter imply heritability less than one; however, the exact value of heritability (the proportion of phenotypic variation explained by genetics) could still vary depending on the extent of genetic variation.
- Mutation rate. It was either set to zero to explore the role of standing genetic variation or varied between 1e-6 and 1e-4 per QTL per gamete. This range covers and exceeds the range of trait-level deleterious mutation rates observed in humans (*57*). Therefore, values at higher end of the range most likely strongly over-estimate the rate of adaptive mutations, which was done deliberately to show that no realistic mutation rate could help sustain genetic variation in the face of strong selection by warming.

The model’s code, available at <u>https://github.com/z0on/Adaptive-pathways-of-coral-populations-on-the-Great-Barrier-Reef</u>, is designed for general modeling of multilocus adaptation in metapopulations. It can read user-supplied files of environmental conditions, population sizes and migration matrices for arbitrary number of populations.

Here, we modeled our five populations with effective population sizes and pairwise migration rates inferred by *dadi*. We modeled identical thermal trends across populations with population-specific offsets. During pre-adaptation period lasting 2000 generations, the temperature was constant on average but experienced random fluctuations across generations drawn from a normal distribution with a standard deviation of 0.25°C (to approximate ENSO events). The temperature was offset by +1.6°C in Wilkie and Magnetic populations and by -1.8°C in the Keppel population, to model differences in midsummer monthly mean temperature among populations (Fig. 1). After 2000 generations a linear increase at 0.05°C per generation was added to simulate warming.

All combinations of parameter settings were run ten times to ensure consistency. We found that with population sizes in thousands, such as in our case, the results were very consistent among independent runs. We therefore did not aggregate results over many replicated runs but show one randomly chosen run for each tested parameter combination.

## Acknowledgements

We wish to thank Ryan Gutenkunst and Benjamin Haller for their continuous support of *dadi* and SLiM users, respectively. The bioinformatics analysis was accomplished using computational resources of the Texas Advanced Computer Center. This study has been supported by NSF (DEB-1054766) grant to M.V.M, ARC (LP120200245) and University of Melbourne ECR grants to E. A.T., a Coral Reef Alliance grant (“Coral Adaptation Challenge”) to E.A.T and M.V.M, Queensland Government funding to L.K.B and AIMS funding to L.K.B. and M.J.V.O

## Data and code availability

The finalized genotyping dataset in VCF format, detailed bioinformatic walkthrough, accessory formatting and plotting scripts, *dadi* scripts and the SLiM model code are available from <u>https://github.com/z0on/Adaptive-pathways-of-coral-populations-on-the-Great-Barrier-Reef</u>. Raw sequencing data has been deposited to National Center for Biotechnology Information’s Short Reads Archive (accession number pending).

## Author contributions

M.V. M. conceived the study, performed genotyping data analysis and all statistical analyses,programmed and analyzed evolutionary simulations, wrote the manuscript and prepared the figures. E. A. T. performed biophysical modeling and contributed to figure preparation and manuscript writing. G. V. A. performed all lab procedures related to enotyping. M. J. H. v. O and L. K. B. provided original coral samples and contributed to manuscript preparation and revision.

## Supplemental Figures

**Figure S1.**
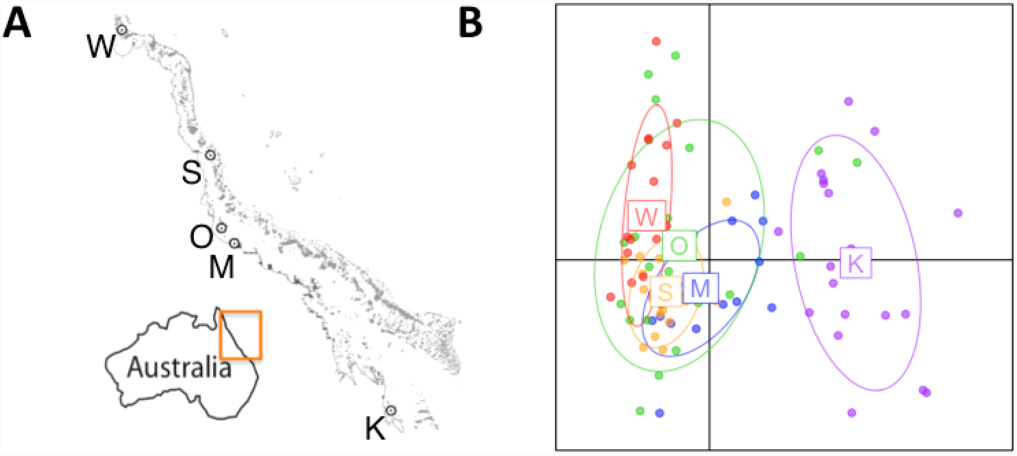
Principal component analysis of genetic diversity in sampled populations. (A) Map of sampled locations with one-letter population identifiers. (B) Principal component analysis of genome-wide genetic variation. On panel D, centroid labels are initial letters of population names as in panel A.

**Figure S2.**
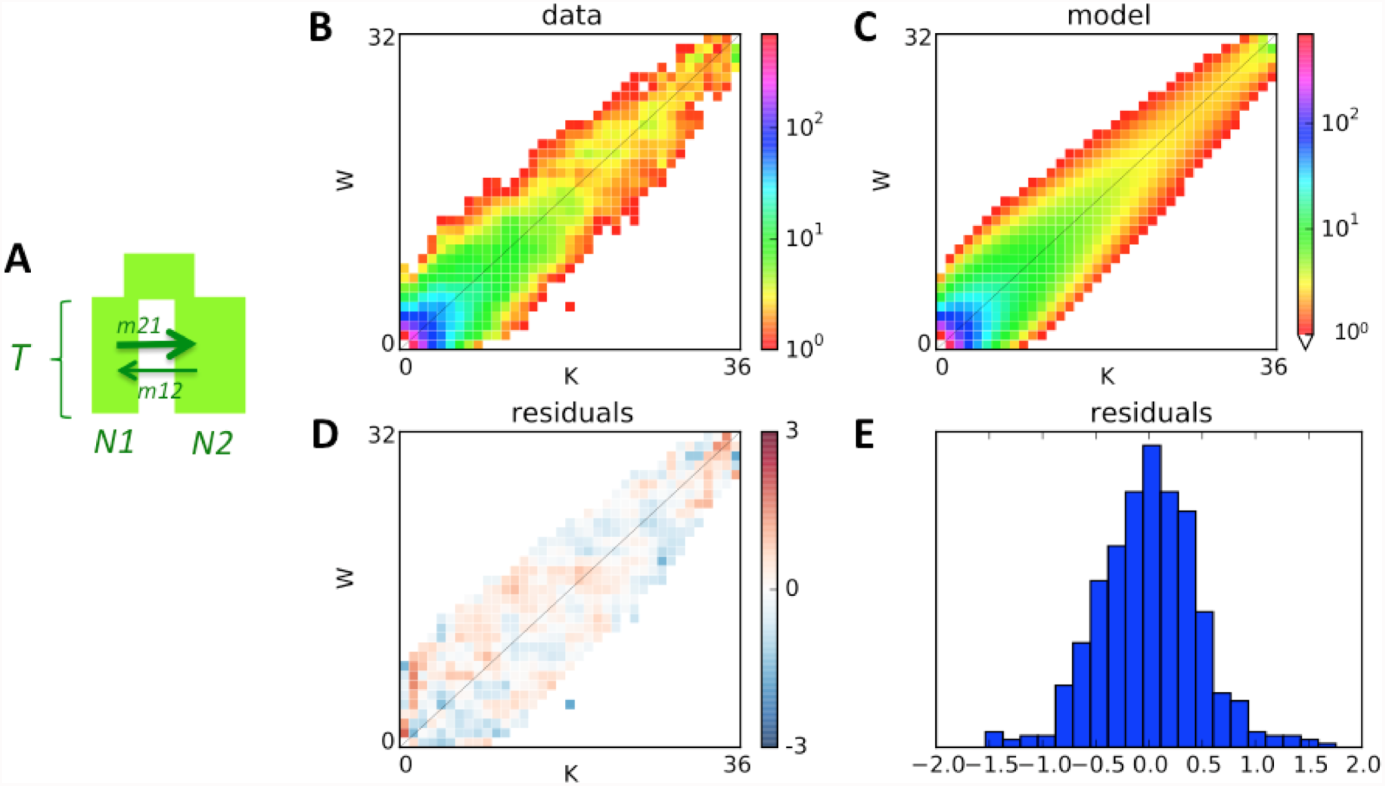
Example of two-population *dadi* model fit. (A) The model: ancestral population splits into two populations of unequal sizes (N1 and N2) some time T in the past, which exchange migrants with different rates depending on direction. (B) Observed allele frequency spectrum comparing Wilkie (W) and Keppel(K) populations. (C) Allele frequency spectrum generated by the fitted model. (D, E) Map and histogram of residuals (absolute scale).

**Figure S3.**
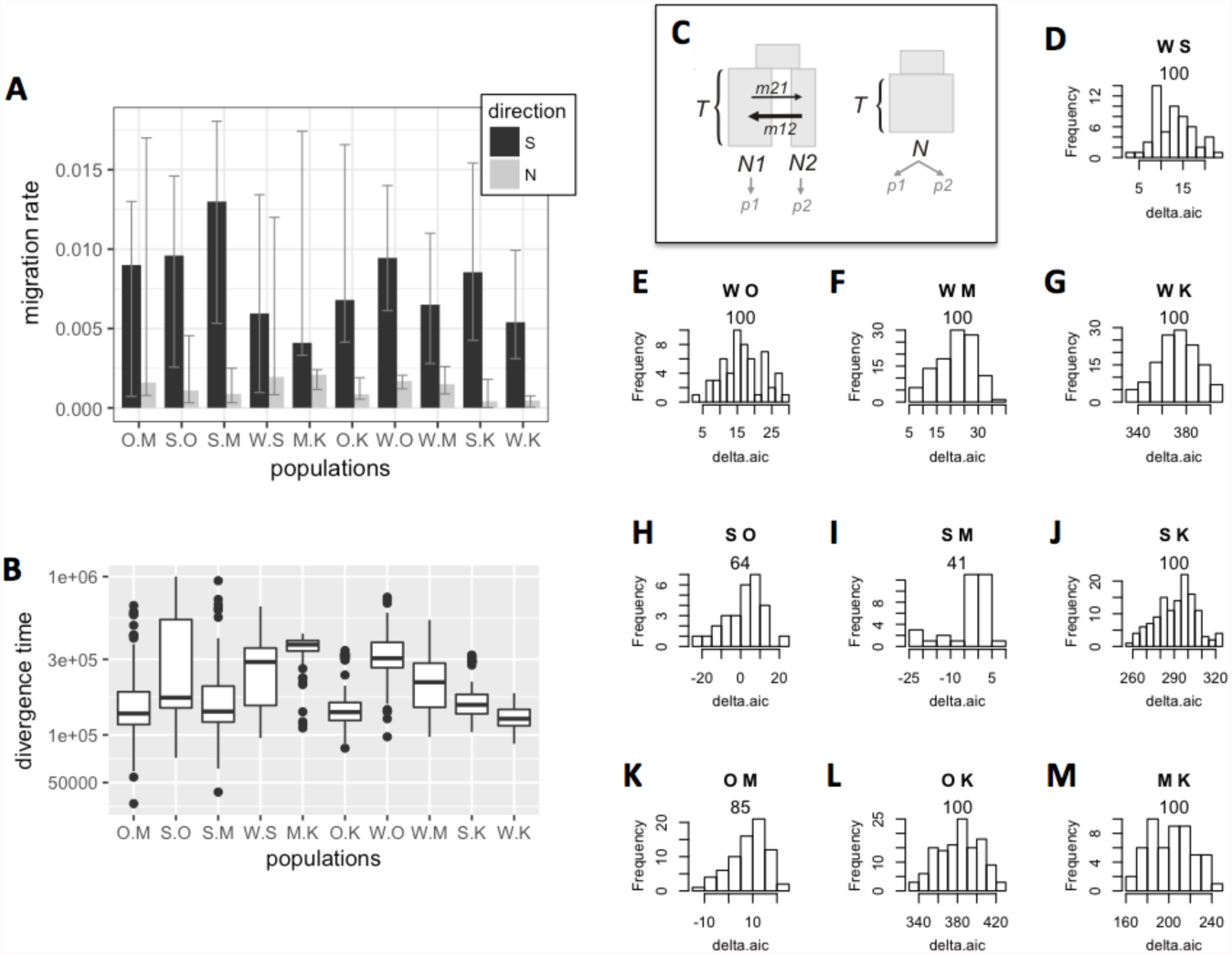
Bootstrap analysis of migration rates, divergence times, and population subdivision. (A)Migration among population pairs, with bootstrap-derived 95% confidence intervals. The pairs are identified on the x-axis and sorted by increasing geographical distance. Black bars - southward migration, grey bars - northward migration. (B) Boxplot of divergence times (in years, y-axis) between pairs of populations (x-axis) across bootstrap replicates. (C) Models being compared: the split model (left) implies populations’ split into two different sizes (N1 and N2) at time T in the past, since when they exchanged migrants at unequal rates depending on direction. No-split model (right) allows for population size change at time T in the past but does not include population split: the two genotyped groups (p1 and p2) are regarded as two samples from the same population. (D-M) Histograms of delta-AIC values comparing split and no-split models (panel A) for bootstrap replicates (bootstrap was performed over genomic contigs of the draft genome of *A. digitifera*). Positive numbers indicate support for the split model. The letters on top of each panel identify compared populations, the number is the proportion of positive bootstrap replicates(i.e., bootstrap support for the full model).

**Figure S4.**
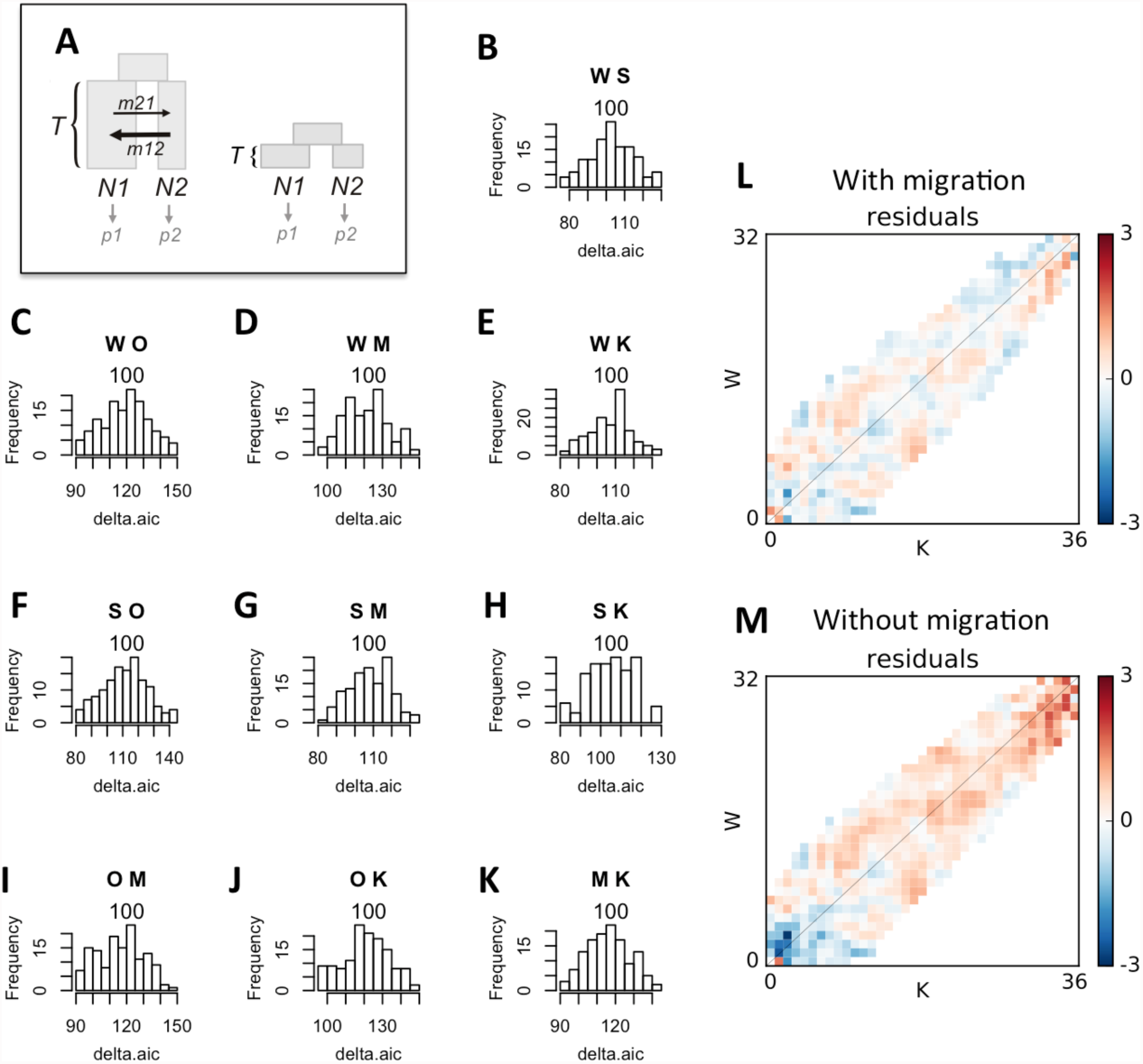
Delta-AIC bootstrap comparison of models with and without migration (A), to confirm that the model with migration and ancient divergence is preferable to the model with no migration but very recent divergence. (B-K) Histograms of delta-AIC values for bootstrap replicates comparing models with and without migration. Positive numbers indicate support for the model with migration. The letters on top of each panel identify compared populations, the number is the proportion of bootstrap replicates supporting the model with migration and ancient split. For all pairs of populations the model of ancient split with migration is strongly supported. (L, M) Example of residuals from the two models. Model without migration under-estimates the number of shared low-frequency polymorphisms and over-estimates the number of shared high-frequency polymorphisms.

**Figure S5.**
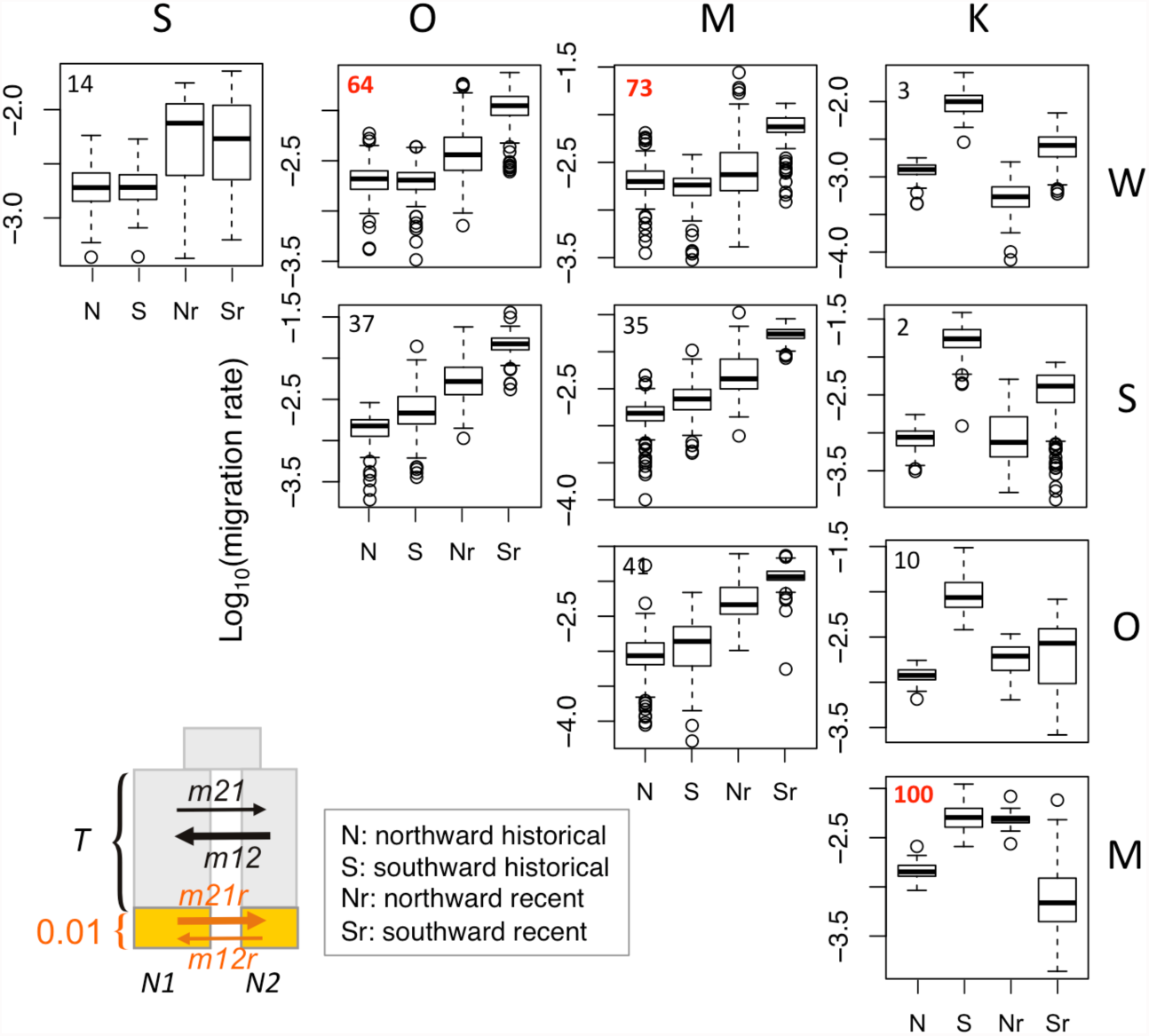
Migration rates inferred by the *dadi* model allowing for the change in migration rates over the last 0.01 T units (15-20 generations or 75-100 years, in our case). Box plots show historical (N, S) and recent (Nr, Sr) migration rates inferred among pairs of population across bootstrap replicates. Numbers in the top left corner are delta-AIC bootstrap support for the model with the recent change in migration compared to the split-with-migration model with no recent change (Fig. 2A, inset). In three cases (OW, MW, and KM) the model with recent migration change is favored. Overall, there is no consistency in recent migration increase from cooler to warmer location; for example, the split model consistently predicts increase in recent immigration to the warmer Magnetic Island (column M).

**Figure S6.**
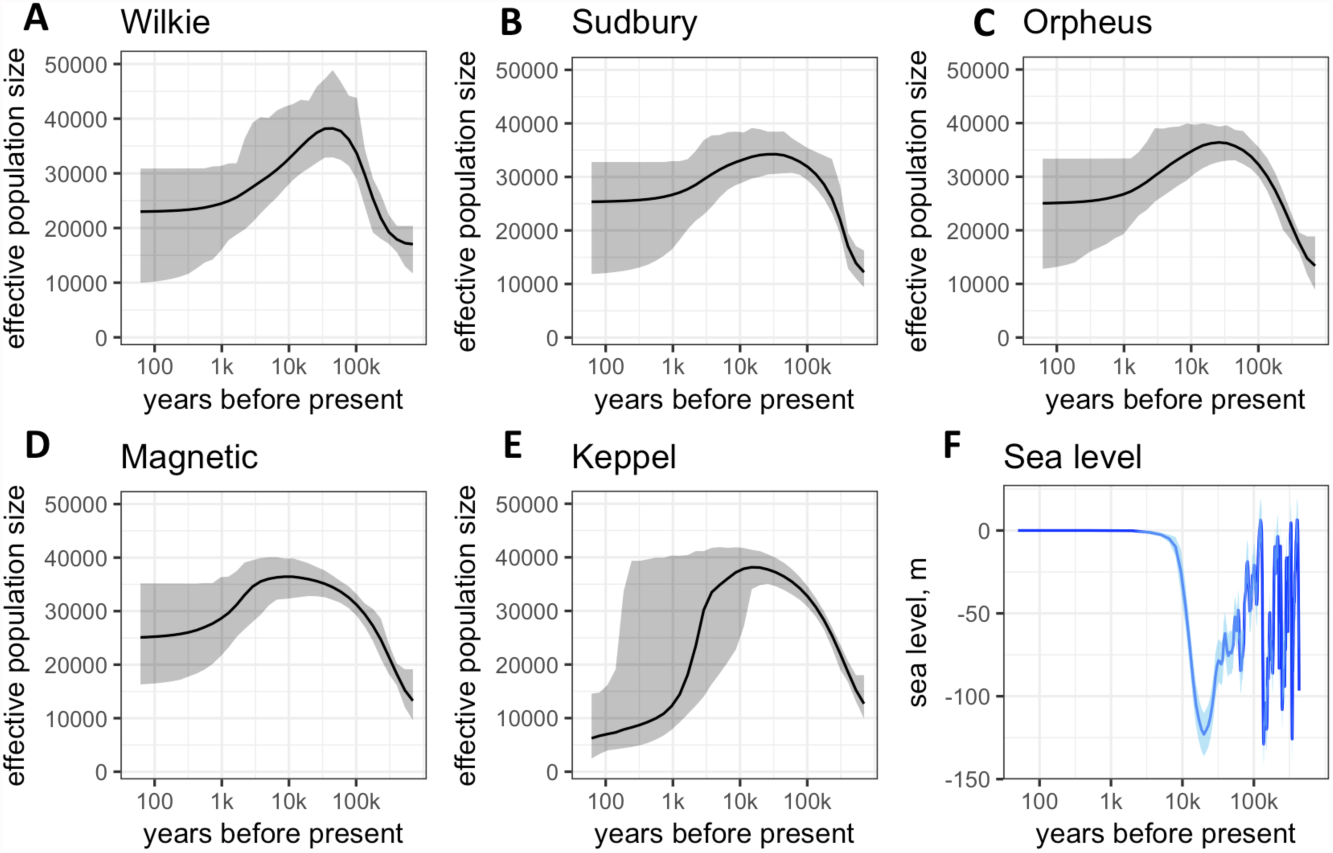
Population history. (A-E) Historical population sizes with bootstrap-derived 95% confidence intervals, according to the two-growth model (Fig. S7 F). (F) Sea level with shaded area corresponding to standard error (*41*).

**Figure S7.**
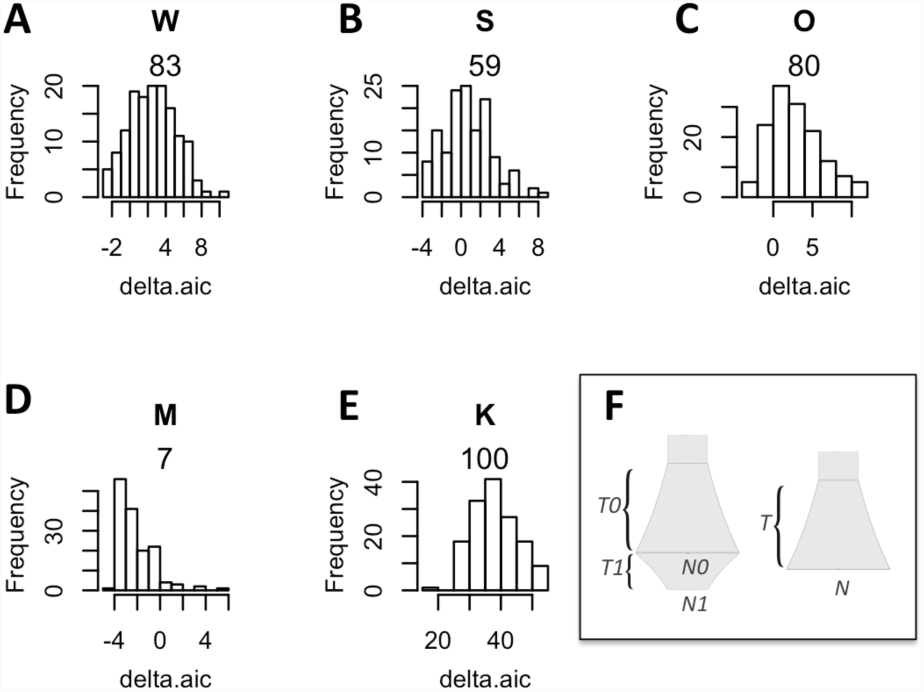
Delta-AIC bootstrap analysis of single-population models. (A-E) Histograms of delta-AIC values for 100 bootstrap replicates comparing two-growths and one-growth models (panel F). Positive numbers indicate support for the two-growth model. The letter on top of each panel identify the population, the number is the proportion of positive bootstrap replicates (i.e., bootstrap support for the two-growth model). The two-growth model is well supported for populations W, O, and K (panels A, C and E), weakly supported for population S (panel B), and not supported for population M (panel D). (F) Models compared. The full model (left) includes two exponential growth periods (any of which could be growth or decline), the reduced model (right) has only one growth period.

**Figure S8.**
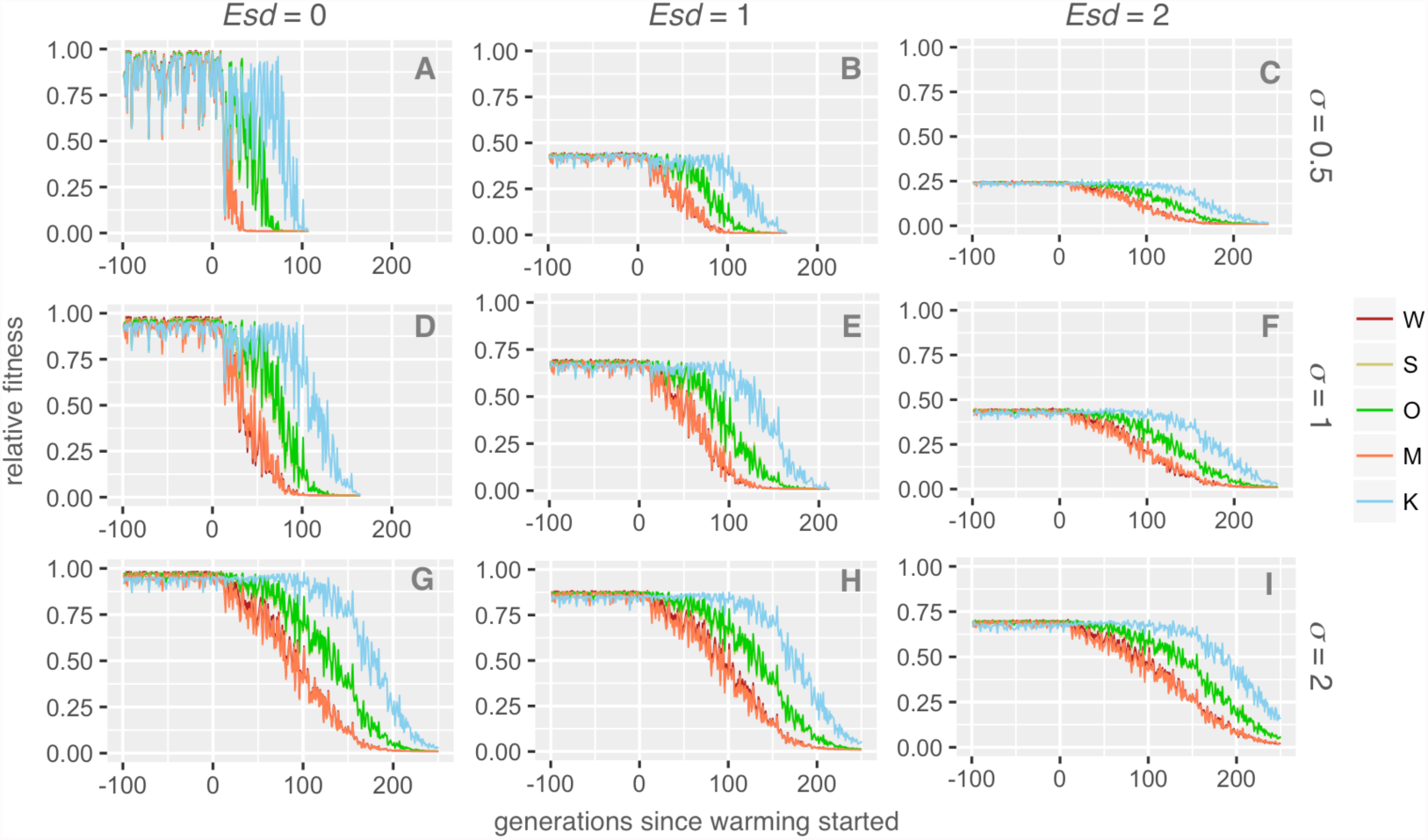
Higher plasticity and lower heritability facilitate metapopulation persistence during warming. The graphs show fitness of populations (relative to maximal fitness at the genetically determined optimum) after pre-adaptation period and under warming, depending on the magnitude of non-heritable component (*Esd*, standard deviation of normally distributed random value added to the sum of QTL effects when calculating individual’s phenotype; higher *Esd* implies lower heritability) and phenotypic plasticity (σ, standard deviation of the Gaussian slope of fitness decline away from the phenotypic optimum, in degrees C). Higher plasticity confers stability against random thermal fluctuations (compare panels A, D and G) and partially rescues the drop in fitness due to high *Esd* (i.e., lower heritability - compare pre-warming generations, from -100 to 0, on panels B, E and H or C, F, and I).

**Figure S9.**
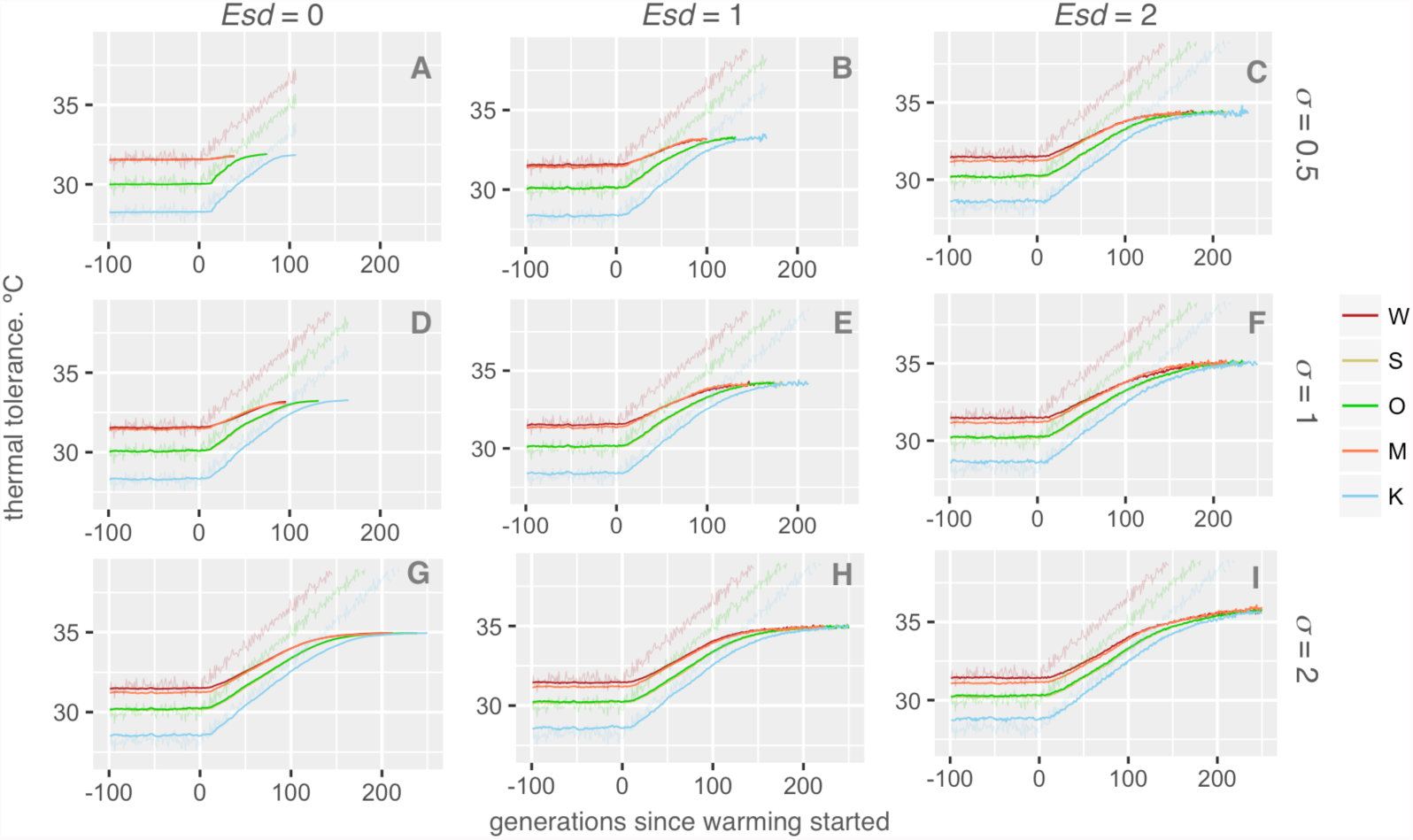
Higher plasticity and lower heritability promote longer and more extensive evolution in response to warming. The graphs show mean thermal tolerance of populations after pre-adaptation period and under warming, depending on the magnitude of non-heritable component (*Esd*, standard deviation of normally distributed random value added to the sum of QTL effects when calculating individual’s phenotype; higher Esd implies lower heritability) and phenotypic plasticity (σ, standard deviation of the Gaussian slope of fitness decline away from the phenotypic optimum, in degrees C).

**Table S1.**
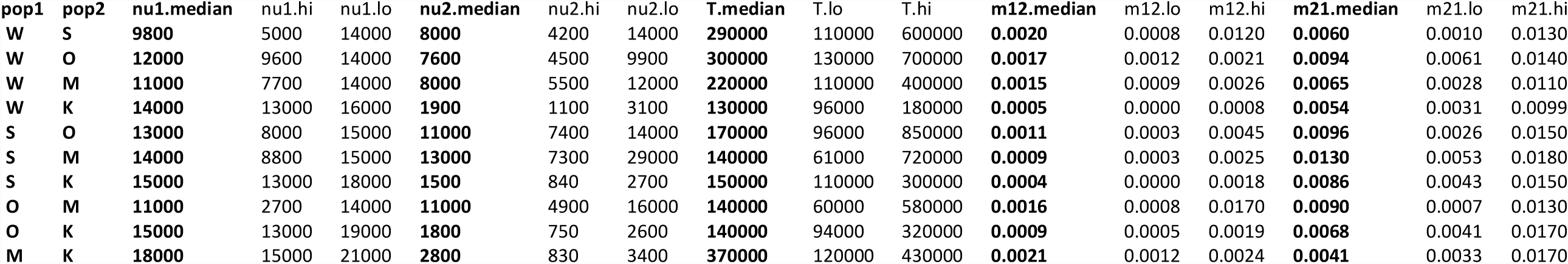
Summary of demographic analysis based on split-with-migration model (Fig. 2 A). “lo” and “hi” values are bootstrap-based 95% confidence limits. This table (S2M_results_summary) as well as full listing of raw parameter values obtained for all bootstrap replicates(s2m_rawParameters) are available at <u>https://github.com/z0on/Adaptive-pathways-of-coral-populations-on-the-Great-Barrier-Reef</u>

## References

1. A. C. Baker, P. W. Glynn, B. Riegl, Climate change and coral reef bleaching: An ecological assessment of long-term impacts, recovery trends and future outlook. Estuar. Coast. Shelf Sci. 80, 435–471 (2008).

2. C. A. Logan, J. P. Dunne, C. M. Eakin, S. D. Donner, Incorporating adaptive responses into future projections of coral bleaching. Glob. Chang. Biol. 20, 125–139 (2014).

3. T. P. Hughes et al., Climate change, human impacts, and the resilience of coral reefs. Science. 301, 929–933 (2003).

4. J. M. Pandolfi, S. R. Connolly, D. J. Marshall, A. L. Cohen, Projecting coral reef futures under global warming and ocean acidification. Science. 333, 418–22 (2011).

5. G. B. B. Dixon et al., Genomic determinants of coral heat tolerance across latitudes. Science. 348, 1460–1462 (2015).

6. J. A. Kleypas et al., Larval connectivity across temperature gradients and its potential effect on heat tolerance in coral populations. Glob. Chang. Biol. 22, 3539–3549 (2016).

7. G. De’ath, K. E. Fabricius, H. Sweatman, M. Puotinen, The 27-year decline of coral cover on the Great Barrier Reef and its causes. Proc. Natl. Acad. Sci. U. S. A. 109, 17995–9 (2012).

8. S. Wang, E. Meyer, J. K. McKay, M. V Matz, 2b-RAD: a simple and flexible method for genome-wide genotyping. Nat. Methods. 9, 808–810 (2012).

9. M. J. H. van Oppen, V. Lukoschek, R. Berkelmans, L. M. Peplow, A. M. Jones, A population genetic assessment of coral recovery on highly disturbed reefs of the Keppel Island archipelago in the southern Great Barrier Reef. PeerJ. 3, e1092 (2015).

10. M. J. H. Van Oppen, L. M. Peplow, S. Kininmonth, R. Berkelmans, Historical and contemporary factors shape the population genetic structure of the broadcast spawning coral, Acropora millepora, on the Great Barrier Reef. Mol. Ecol. 20, 4899–4914 (2011).

11. R. N. Gutenkunst, R. D. Hernandez, S. H. Williamson, C. D. Bustamante, Inferring the joint demographic history of multiple populations from multidimensional SNP frequency data. PLoS Genet. 5, e1000695 (2009).

12. D. J. Ayre, T. P. Hughes, Genotypic diversity and gene flow in brooding and spawning corals along the Great Barrier Reef, Australia. Evolution (N. Y). 54, 1590–1605 (2000).

13. K. J. Miller, D. J. Ayre, The role of sexual and asexual reproduction in structuring high latitude populations of the reef coral Pocillopora damicornis. Heredity (Edinb). 92, 557–568 (2004).

14. M. C. Whitlock, D. E. McCauley, Indirect measures of gene flow and migration:FST≠1/(4Nm+1). Heredity (Edinb). 82, 117–125 (1999).

15. J. D. Robinson, A. J. Coffman, M. J. Hickerson, R. N. Gutenkunst, Sampling strategies for frequency spectrum-based population genomic inference. Bmc Evol. Biol. 14, 254 (2014).

16. E. A. Treml, J. Roberts, P. N. Halpin, H. P. Possingham, C. Riginos, The emergent geography of biophysical dispersal barriers across the Indo-West Pacific. Divers. Distrib. 21, 465–476 (2015).

17. E. A. Treml et al., Reproductive Output and Duration of the Pelagic Larval Stage Determine Seascape-Wide Connectivity of Marine Populations. Integr. Comp. Biol. 52, 525–537 (2012).

18. J. Lough, A. Hobday, Observed climate change in Australian marine and freshwater environments. Mar. Freshw. Res. 62, 984–999 (2011).

19. B. Charlesworth, Fundamental concepts in genetics: Effective population size and patterns of molecular evolution and variation. Nat. Rev. Genet. 10, 195–205 (2009).

20. W. Renema et al., Are coral reefs victims of their own past success? Sci. Adv. 2, e1500850 (2016).

21. A. Keinan, A. G. Clark, Recent Explosive Human Population Growth Has Resulted in an Excess of Rare Genetic Variants. Science. 336, 740–743 (2012).

22. B. C. Haller, P. W. Messer, SLiM 2: Flexible, interactive forward genetic simulations. Mol. Biol. Evol. 34, 230–240 (2017).

23. M. Vanessa, B. Baria, D. W. Dela Cruz, R. D. Villanueva, J. R. Guest, Spawning of three-year-old Acropora millepora corals reared from larvae in Northern Philippines. Bull. Mar. Sci. 88, 61–62(2012).

24. IPCC, Climate Change 2007 - The Physical Science Basis. Contribution of Working Group I to the Fourth Assessment Report of the IPCC (Cambridge University Press, New York, NY, 2007).

25. R. Berkelmans, M. J. H. van Oppen, The role of zooxanthellae in the thermal tolerance of corals: a“nugget of hope” for coral reefs in an era of climate change. Proc. R. Soc. B-Biological Sci. 273, 2305–2312 (2006).

26. A. H. Baird, J. R. Guest, B. L. Willis, Systematic and biogeographical patterns in the reproductive biology of scleractinian corals. Annu. Rev. Ecol. Evol. Syst. 40, 551–571 (2009).

27. K. Quigley, B. Willis, L. Bay, Heritability of the Symbiodinium community in vertically- and horizontally-transmitting broadcast spawning corals. bioRxiv. **doi: https** (2017).

28. E. J. Howells et al., Coral thermal tolerance shaped by local adaptation of photosymbionts. Nat. Clim. Chang. 2 (2011), pp. 116–120.

29. F. P. Palstra, D. J. Fraser, Effective/census population size ratio estimation: a compendium and appraisal. Ecol. Evol. 2, 2357–2365 (2012).

30. S. R. Palumbi, D. J. Barshis, N. Traylor-Knowles, R. A. Bay, Mechanisms of reef coral resistance to future climate change. Science. 344, 895–8 (2014).

31. E. J. Howells, R. Berkelmans, M. J. H. van Oppen, B. L. Willis, L. K. Bay, Historical thermal regimes define limits to coral acclimatization. Ecology. 94, 1078–1088 (2013).

32. D. Nettle, M. Bateson, Adaptive developmental plasticity: what is it, how can we recognize it and when can it evolve? Proc. R. Soc. B Biol. Sci. 282, 20151005 (2015).

33. Z. T. Richards, D. J. Miller, C. C. Wallace, Molecular phylogenetics of geographically restricted Acropora species: Implications for threatened species conservation. Mol. Phylogenet. Evol. 69,837–851 (2013).

34. M. Lynch, Evolution of the mutation rate. Trends Genet. 26, 345–52 (2010).

35. T. J. Done, Phase shifts in coral reef communities and their ecological significance. Hydrobiologia. 247, 121–132 (1992).

36. O. Hoegh-Guldberg et al., Assisted colonization and rapid climate change. Science. 321, 345–346(2008).

37. S. N. Aitken, M. C. Whitlock, Assisted Gene Flow to Facilitate Local Adaptation to Climate Change. Annu. Rev. Ecol. Evol. Syst. 44, 367–388 (2013).

38. Great Barrier Reef Marine Park Authority, Interim report on the environmental impacts of the 2016 coral bleaching event.

39. G. B. Dixon, L. K. Bay, M. V. Matz, Bimodal signatures of germline methylation are linked with gene expression plasticity in the coral Acropora millepora. BMC Genomics. 15, 1109 (2014).

40. C. Shinzato et al., Using the Acropora digitifera genome to understand coral responses to environmental change. Nature. 476, 320–U82 (2011).

41. M. J. H. van Oppen, B. J. McDonald, B. Willis, D. J. Miller, The Evolutionary History of the Coral Genus Acropora (Scleractinia, Cnidaria) Based on a Mitochondrial and a Nuclear Marker: Reticulation, Incomplete Lineage Sorting, or Morphological Convergence? Mol. Biol. Evol. 18, 1315–1329 (2001).

42. B. F. Voight, S. Kudaravalli, X. Wen, J. K. Pritchard, A map of recent positive selection in the human genome. PLoS Biol. 4, e72 (2006).

43. I. K. Jordan et al., A universal trend of amino acid gain and loss in protein evolution. Nature. 433,633–8 (2005).

44. A. Mckenna et al., The Genome Analysis Toolkit: A MapReduce framework for analyzing next-generation DNA sequencing data. Genome Res. 20, 1297–1303 (2010).

45. P. Danecek et al., The variant call format and VCFtools. Bioinformatics. 27, 2156–8 (2011).

46. T. Jombart, adegenet: a R package for the multivariate analysis of genetic markers. Bioinformatics.24, 1403–5 (2008).

47. D. H. Alexander, J. Novembre, K. Lange, Fast model-based estimation of ancestry in unrelated individuals. Genome Res. 19, 1655–64 (2009).

48. T. Günther, G. Coop, Robust identification of local adaptation from allele frequencies. Genetics.195, 205–20 (2013).

49. M. Tine et al., European sea bass genome and its variation provide insights into adaptation to euryhalinity and speciation. Nat. Commun. 5, 5770 (2014).

50. E. A. Treml, P. N. Halpin, D. L. Urban, L. F. Pratson, Modeling population connectivity by ocean currents, a graph-theoretic approach for marine conservation. Landsc. Ecol. 23, 19–36 (2008).

51. E. P. Chassignet et al., The HYCOM (HYbrid Coordinate Ocean Model) data assimilative system.J. Mar. Syst. 65, 60–83 (2007).

52. R. C. Babcock et al., Synchronous Spawnings of 105 Scleractinian Coral Species on the Great-Barrier-Reef. Mar. Biol. 90, 379–394 (1986).

53. S. W. Davies, E. A. Treml, C. D. Kenkel, M. V. Matz, Exploring the role of Micronesian islands in the maintenance of coral genetic diversity in the Pacific Ocean. Mol. Ecol. 24, 70–82 (2015).

54. S. R. Connolly, A. H. Baird, Estimating dispersal potential for marine larvae: dynamic models applied to scleractinian corals. Ecology. 91, 3572–3583 (2010).

55. P. K. Smolarkiewicz, J. Szmelter, An MPDATA-based solver for compressible flows. Int. J. Numer. Methods Fluids. 56, 1529–1534 (2008).

56. R. K. Cowen, G. Gawarkiewicz, J. Pineda, S. Thorrold, F. Werner, Population Connectivity in Marine Systems: An Overview. Oceanography. 20, 14–21 (2007).

57. M. R. Cummings, Human heredity: principles and issues (Brooks/Cole, Cengage Learning, 2014).

